# The flickering connectivity system of the north Andean páramos

**DOI:** 10.1101/569681

**Authors:** Suzette G.A. Flantua, Aaron O’Dea, Renske E. Onstein, Henry Hooghiemstra

## Abstract

**Aim:** To quantify the effect of Pleistocene climate fluctuations on habitat connectivity across páramos in the Neotropics.

**Location:** The Northern Andes

**Methods:** The unique páramos habitat underwent dynamic shifts in elevation in response to changing climate conditions during the Pleistocene. The lower boundary of the páramos is defined by the upper forest line, which is known to be highly responsive to temperature. Here we reconstruct the extent and connectivity of páramos over the last 1 million years (Myr) by reconstructing the UFL from the long fossil pollen record of Funza09, Colombia, and applying it to spatial mapping on modern topographies across the Northern Andes for 752 time slices. Data provide an estimate of how often and for how long different elevations were occupied by páramos and estimates their connectivity to provide insights into the role of topography in biogeographic patterns of páramos.

**Results:** Our findings show that connectivity amongst páramos of the Northern Andes was highly dynamic, both within and across mountain ranges. Connectivity amongst páramos peaked during extreme glacial periods but intermediate cool stadials and mild interstadials dominated the climate system. These variable degrees of connectivity through time result in what we term the ‘flickering connectivity system’. We provide a visualization (video) to showcase this phenomenon. Patterns of connectivity in the Northern Andes contradict patterns observed in other mountain ranges of differing topographies.

**Main conclusions:** Pleistocene climate change was the driver of significant elevational and spatial shifts in páramos causing dynamic changes in habitat connectivity across and within all mountain ranges. Some generalities emerge, including the fact that connectivity was greatest during the most ephemeral of times. However, the timing, duration and degree of connectivity varied substantially among mountain ranges depending on their topographic configuration. The flickering connectivity system of the páramos uncovers the dynamic settings in which evolutionary radiations shaped the most diverse alpine biome on Earth.

## 1. INTRODUCTION

Mountains are regarded as powerhouses of biodiversity in the world (Barthlott, Rafiqpoor, Kier, & Kreft, 2005; Kreft & Jetz, 2007; Antonelli et al., 2018) and harbour numerous examples of very rapid and recent species diversifications (‘radiations’; Hughes & Atchison, 2015). It is thought that a large part of this diversity arose geologically recently, during the Plio-Pleistocene (last 5.3 million years, [Ma]), but there is no consensus on the drivers of these radiations. One favoured hypothesis is that the combination of high topographic relief and Plio-Pleistocene climatic oscillations led to rapidly changing distributions of montane species, which generated new lineages (e.g. Qian & Ricklefs, 2000; Graham et al., 2014; Mutke, Jacobs, Meyers, Henning, & Weigend, 2014). However, the relative contributions of isolation (e.g. Schönswetter, Stehlik, Holderegger, & Tribsch, 2005; Wallis, Waters, Upton, & Craw, 2016; Weir, Haddrath, Robertson, Colbourne, & Baker, 2016) vs. gene flow and dispersal (e.g. Smith et al., 2014; Cadena, Pedraza, & Brumfield, 2016; Kolář, Dušková, & Sklenář, 2016; Knowles & Massatti, 2017) in driving fast diversification rates (i.e. the ‘species-pump’ effect, Rull, 2005; Rull & Nogué, 2007; Winkworth, Wagstaff, Glenny, & Lockhart, 2005; Ramírez-Barahona & Eguiarte, 2013; Steinbauer et al., 2016; Flantua & Hooghiemstra, 2018) are still debated. It is likely that these radiations have been the results of the interchange between phases of isolation, causing allopatric, *in situ* speciation, and connectivity, triggering diversification through dispersal and settlement in new areas and hybridization of differentiated taxa from previously isolated populations (Flantua & Hooghiemstra, 2018). The fastest and most spectacular radiations may therefore occur in mountain regions with variable degrees of past connectivity and isolation during climate fluctuations, which, complex in space and time, are inherently related to the mountain topography (Flantua & Hooghiemstra, 2018). It is therefore critical to quantify connectivity of montane habitats using our understanding of topography and past climate fluctuations (**Fig. 1**).

**Figure 1.**
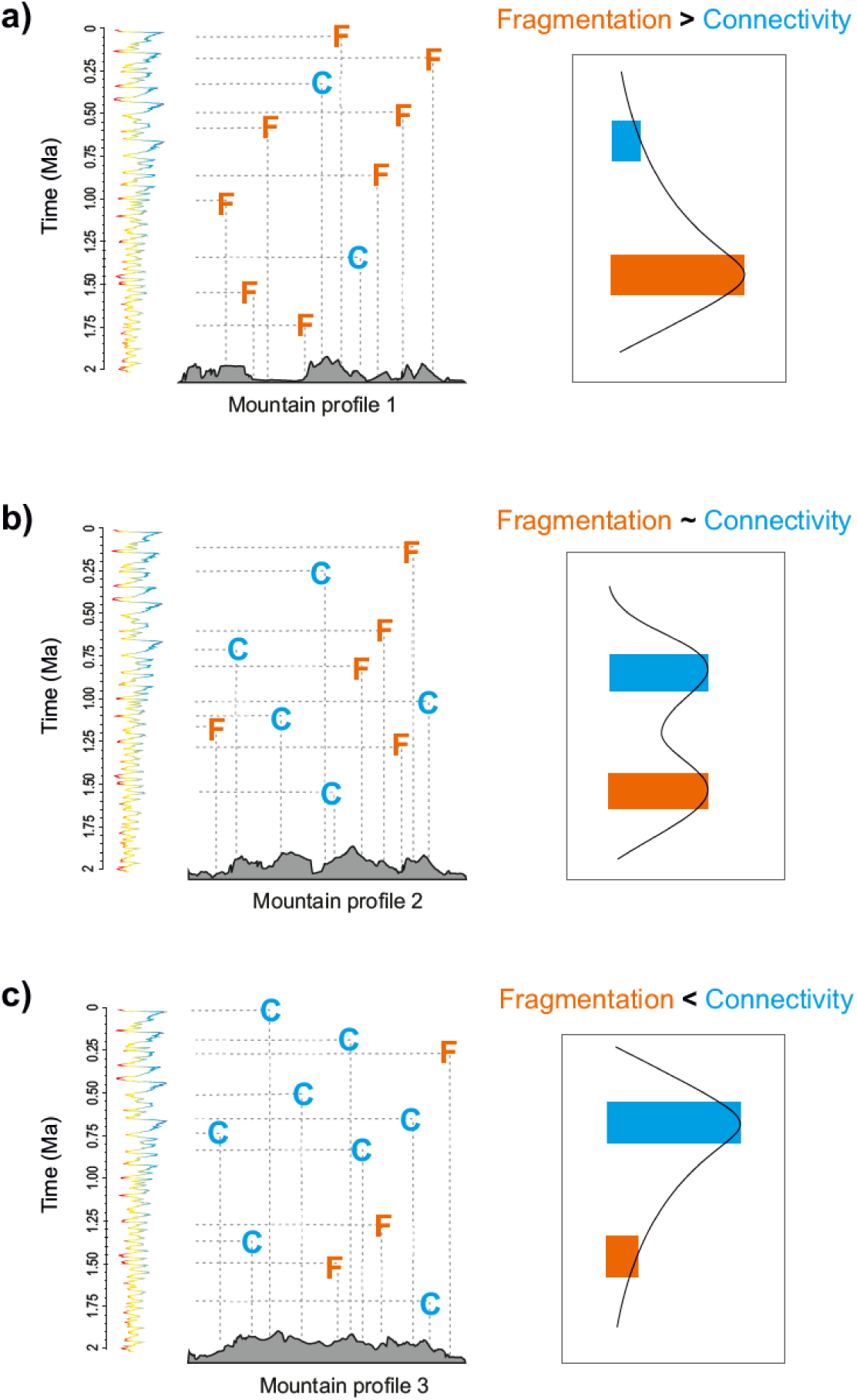
Connectivity and fragmentation in a mountain landscape. Connectivity (blue) and fragmentation (orange) events ocurred in a spatially and temporally variable manner. This complex pattern in space (latitude, longitude, elevation) and time resemblances a multi-dimensional “mountain fingerprint” which is unique for each mountain range (Flantua & Hooghiemstra, 2018). Three hypothetical mountain profiles are shown with elevational shifts in vegetation distribution driven by climate change (pollen-based record at the left indicating temperature). We recognise mountains with where a) only few events of connectivity occurred during the Pleistocene compared to fragmentation events (‘fragmentation-prone mountain fingerprint’), connectivity events interchanged with isolation events in an evenly manner (‘mixed connectivity-fragmentation mountain fingerprint’), c) connectivity is facilitated and occured more often than fragmentation events (‘connectivity-prone mountain fingerprint’). The right panel is only based on frequency, not the duration of each event.

The Northern Andes is an ideal model system to quantify connectivity, due to the large variation in topography and the advanced paleoecological knowledge on Plio-Pleistocene climate fluctuations derived during the last five decades (Hooghiemstra & Flantua, 2019). The Northern Andes is topographically-rich with high elevations, steep ridges and valleys (see illustrations by Von Humboldt during his trips in Latin America, 1773-1858), composed of several mountain ranges, some of which are parallel running from North to South. The area hosts the treeless tundra-like alpine biome, the páramos, regarded the richest alpine flora in the world in terms of endemism and species richness (Sklenář, Hedberg, & Cleef, 2014) and is known for its bursts of Plio-Pleistocene species diversification amongst plants (see overview in Hughes & Atchison, 2015). In terms of quantifying Plio-Pleistocene temperature fluctuations, the palaeoecological history of the páramos has been studied extensively (e.g. Van der Hammen, 1974; Cleef, 1979; Hooghiemstra, 1984; Hooghiemstra & Van der Hammen, 2004) because of the unique high elevation fossil pollen records that cover most of the Pleistocene (Groot et al., 2011; Groot, Hooghiemstra, Berrio, & Giraldo, 2013; Bogotá-Angel et al., 2011; Bogotá-A., Hooghiemstra, & Berrio, 2016; Torres, Hooghiemstra, Lourens, & Tzedakis, 2013). Under current conditions, the páramos form isolated archipelagos of alpine (sky) islands (McCormack, Huang, & Knowles, 2009) but the rich collection of fossil pollen sequences throughout this region (Flantua et al., 2015) show that the páramos underwent substantial elevational shifts during the Pleistocene, resulting in extensive changes in surface area and connectivity (Van der Hammen, 1974; Hooghiemstra & Van der Hammen, 2004; Flantua et al., 2014; Sklenář et al., 2014). Thus, the topographic diversity and the robust catalogue of palaeoecological reconstructions make the Northern Andes a highly suitable model region to explore patterns of connectivity in mountain biomes in response to Pleistocene climate fluctuations.

In this study, we aim to quantify the biogeographic changes of the páramos in terms of spatial scale and connectivity based on modern topography and pollen-based records of past climate change. Specifically, we developed a novel tool to explore the complex temporal and spatial patterns of páramo connectivity. We constrain our model by using the last 1 Myr of the high-resolution fossil pollen record of Funza09, a 586 m deep core taken from the Bogotá basin of Colombia (Torres et al., 2013). Available surface area (Elsen & Tingley, 2015) and connectivity (Flantua et al., 2014; Bertuzzo et al., 2016) is variable along elevational gradients of mountains. We therefore hypothesize that the different mountain ranges that compose the Northern Andes display variable patterns of past páramo connectivity dependent upon their topography (**Fig. 1**). We discuss the implications of our outcomes for evolutionary processes and how defining and quantifying past connectivity in mountain systems is essential to help reveal mechanisms of ecological, biogeographical and evolutionary change. Ultimately, our quantification of páramo connectivity through space and time provides a unique opportunity to disentangle some of the mechanistic drivers (‘modulators’) of radiations in this biome (Bouchenak-Khelladi, Onstein, Xing, Schwery, & Linder, 2015).

## 2. METHODS

### 2.1 Geographical features

The Northern Andes (ca. 448.000 km^2^) covers parts of Venezuela, Colombia and Ecuador (**Fig. 2a**), and can be partitioned into six principal mountain ranges or ‘cordilleras’ (**Fig. 2c**), namely the Sierra Nevada de Santa Marta (SNSM), Cordillera de Mérida, Eastern, Central and Western Cordillera and the Ecuadorian Cordilleras. Most of the Northern Andes is considered a highly to extremely high rugged landscape (**Fig. 2b**; See mountain illustrations by Von Humboldt (1845)where the high peaks and deep inter-Andean valleys cause strong contrasts in climate throughout the region (Flantua et al., 2016). Surface area in mountains does not decrease monotonically with elevation as has been shown previously in southern Colombia by Flantua et al. (2014) and on a global scale by Elsen & Tingley (2015). The Northern Andes shows a decrease of surface area going upslope where there is a slight peak around 900-1200 m asl but then continues to decrease up to 6260 m asl (**Fig. 2d**), following a typical ‘pyramid shape’. However, the different cordilleras show different patterns of elevational surface area (Fig.2d) where the Eastern Cordillera shows a sharp peak around 2600 m asl and the Ecuadorian Cordillera shows high values of surface area at much higher elevations than the other cordilleras (**for more details see Table S1.1, Appendix S1 in Supporting Information**). Of all tropical alpine floras, such as in East Africa and New Guinea, the páramos are home to the highest species richness and endemism (Luteyn, 1999; Sklenář, Dušková, & Balslev, 2011), with low between-mountain similarity in species; (Sklenář et al., 2014). The páramos today are spread out over the Northern Andes as an archipelago of small and highly fragmented páramo complexes (**Figure S2.1**, **Appendix S2**).

**Figure 2.**
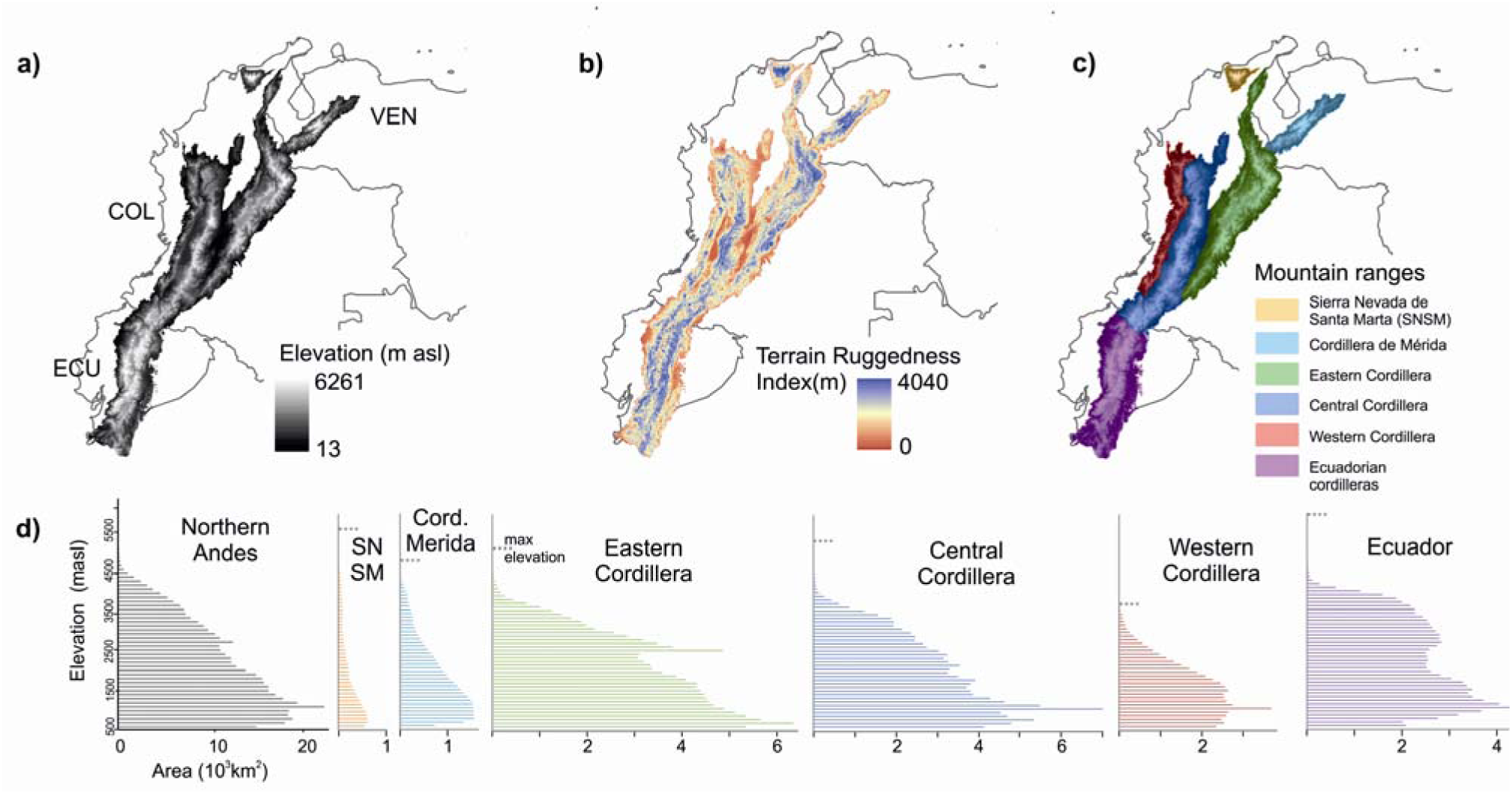
Hypsographic curves of the Northern Andes. a) Elevation (m asl). b) Terrain ruggedness index calculates the sum change in elevation between a grid cell and its eight neighbor grid cells (Riley et al. (1999) using a ca. 30m DEM (NASA STRM Global 1arc second V003). c) Delimitation of mountain ranges. d) Elevational availability of surface area for the Northern Andes and each mountain range separately shown for 100 m bins. Hypsographic curves based on the Shuttle Radar Topography Mission 1-arc second Digital Terrain Elevation Data (~ 30 m resolution; USGS), taking an elevational threshold of 500 m asl as the horizontal reference plane. Maximum elevation per cordillera is indicated. VEN: Venezuela; COL: Colombia; ECU: Ecuador.

### 2.2 Quantifying temperature and upper forest line based on fossil pollen data

To quantify temperature fluctuations during the Pleistocene (and consequently estimate páramo connectivity), we used fossil pollen data from the Northern Andes. The composite pollen record Funza09 (4.83°N, 75.2°W; 2550 m asl, **Fig. S2.1. Red star**) reveals vegetation and climate dynamics over the past 2.25 Myr (Torres et al., 2013). We reconstructed the páramos’ elevational fluctuations, and consequently páramo connectivity, by estimating the upper forest line (the transition from the upper montane forest to the páramos; UFL) from the Funza09 record. Though this record covers the last 2.2 Myr, we only used the last 1 Myr as this interval reflects continuous lake conditions in comparison with variable hydrological conditions between 2.2-1 Ma which obscure the quantification of changes to the UFL. We follow the methodology described and implemented by Hooghiemstra (1984), Groot et al. (2011), and Hooghiemstra et al. (2012) to derive the Andean UFL and paleotemperature curve (for detailed methodology on the UFL reconstruction see **Appendix S3**)

### 2.3 Calculations of connectivity per páramo “island”

To calculate the degree of connectivity between páramos, we used a graph-based habitat availability index called probability of connectivity (PC) metric. This metric takes into account the area of the páramo “island” itself and the distances to other islands where a user-defined distance threshold defines the ‘reachability’ of other islands (Saura & Pascual-Hortal, 2007; Saura, Estreguil, Mouton, & Rodríguez-Freire, 2011), even if they are not physically connected (i.e. ‘structural connectivity’, Tischendorf & Fahrig, 2000). The metric assigns a value to each páramo island representing its contribution in maintaining the overall connectivity of the páramo biome (Saura & Pascual-Hortal, 2007; Saura et al., 2011). The total PC is built up in three ‘fractions’, namely the ‘intrapatch’, the ‘flux’, and the ‘connector’ fractions (Saura & Rubio, 2010). The first fraction focusses on the available surface area and habitat quality (if applicable) within the individual island. The second fraction assesses how well the individual island is connected to other islands given additional importance to the other islands’ attributes (surface and quality) and its strategic position to other páramo islands. The third fraction quantifies the contribution of the island to maintain connectivity between the rest of the islands, in other words its role as an intermediate stepping stone between non-adjacent islands. Additionally, we calculated the equivalent connected area (ECA), which is derived directly from the PC, as a measure of the overall connectivity of a region (Saura et al., 2011). Conefor Sensinode 2.2 software and ESRI ArcGIS 10.3 were used to calculate the straight-line distances between islands, the PC and ECA (Saura & Pascual-Hortal, 2007; Saura & Torné, 2009). We calculated connectivity for the entire Northern Andes and for each mountain range separately.

### 2.4 Calculations of corridors between páramo islands

We identified corridors between páramo islands within and between cordilleras under different climatic conditions. We used the Gnarly Landscape Utilities (V0.1.3; McRae, Shirk, & Platt, 2013) with ESRI ArcGIS 10.3 to create a raster grid of ‘landscape resistance’ based on ruggedness (**Fig. 2b**) and habitat suitability. We assumed an increased landscape resistance with increased ruggedness, assigning values between 0 (no resistance) to 100 (maximum resistance) using an equal interval classification. For the habitat suitability map, we started by assigning a “perfectly suitable” score of 100 to each páramo island, while outside the island the score of 0 reflects maximum unsuitability. To soften this boundary, an exponential decay function was then used by increasing resistance in 5 elevational steps of 100 m where we assigned a suitability score of 40 to the boundary of the páramo. As a result of the decay function the highest suitability of páramo - its core area - was restrained 200 m above the UFL and 200 m below the snowline.

We used Linkage mapper to calculate the least-cost pathways, or corridors, based on the produced raster grid of landscape resistance (McRae & Kavanagh, 2011). These corridors are expressed as ‘conductance maps’ that represent gradients of cumulative corridors. Where the densities of corridors is highest, it is assumed that there is a high probability of dispersal and migration possible between islands (McRae, Dickson, Keitt, & Shah, 2008). The full landscape of the Northern Andes is considered an area where corridors could exist, with exception of the region between SNSM and the Sierra de Perijá (**Fig. S2.1**).

We resampled the 30 m Digital Elevation Model (DEM, **Fig. 2**) to a 1 km resolution to reduce computing time for each Linkage mapper down to on average 2 hours. We allowed Linkage mapper to create corridors through (instead of only between) core areas to represent the full arsenal of connectivity through the landscape. Only corridors between páramo islands larger than 1 km^2^ were considered at any given moment in time. From the final output maps, only values lower than 200k conductance (default threshold) are selected to highlight the strongest corridors. The outputs were weighted according to the percentage of time they occurred during the last 1 Myr.

## 3. RESULTS

### 3.1 A million years of temperature fluctuations

Temperatures at Funza (2550 masl) are estimated to have fluctuated between ca. 15 and 6°C causing an estimated maximum 1600 m elevational shift of the UFL between ca. 3500 and ca. 1900 m asl (**Fig. 3**). The Pleistocene glacial-interglacial dynamics were not replicated cycles of temperature change showing repeated patterns of high and lows, but display a high temporal variability between each glacial-interglacial cycle. Conditions similar to the current warm, interglacial conditions occurred several times during the last 1 Myr and accounted for around a quarter of the time. Extreme cool glacial conditions, ~ 6 - 8°C cooler than today, were relatively rare, occurring less than 10 percent of the time. On the whole, intermediate cool stadials and mild interstadials dominated the last 1 Myr, occurring over two thirds of the time.

**Figure 3.**
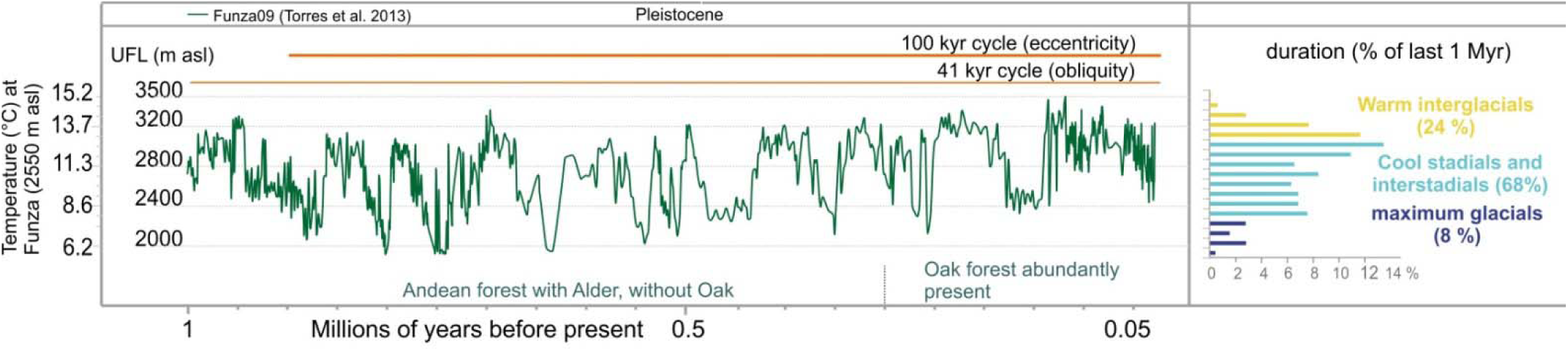
Upper forest line (UFL) curve of Funza09 (Torres et al. 2013) and reconstructed temperature record covering the last 1 Myr (last ca. 30 kyr BP not included).

### 3.2 Calculations of páramo connectivity

Our estimations on the spatial and elevational extent of ancient páramos and their connectedness at different times in the past reveals that páramos underwent frequent spatial alterations between fragmented and connected spatial configurations, but the exact patterns were highly dependent on mountain chain topography (**Fig. 4a,b. See Appendices 4 and 5**). The páramos in the Ecuadorian Cordillera generally maintained a high degree of connectivity over the last 1 Myr, rarely enduring severe fragmentation. Fragmentation did however occur when the snowline plunged significantly during colder and wetter glacial periods, causing a break up of páramo areas on lateral flanks of the mountains. Likewise, the level of connectivity between páramos on the Central Cordillera fragmented substantially through a descending snowline, breaking the upper elevation limit of páramo connectivity. In contrast, the Eastern Cordillera shifted substantially between periods of connectivity and fragmentation, always, however, maintaining two large páramo islands surrounded by smaller ‘satellite islands’. Páramos in the Cordillera de Mérida seem to have been restricted during interglacials to one core area only, while during colder periods a relatively high fragmentation is observed possibly due to glaciers pushing páramos to lateral distributions. Here connectivity increased mainly towards the southwest and during colder periods (UFL ≤ 2300 m asl). The páramos of the SNSM and the Western Cordillera endured the highest degree of rates of change in fragmentation of all ranges. In the latter, páramo habitats are estimated to have often completely disappeared. In contrast, páramos of the Central Cordillera maintained a long latitudinal distribution, forming a chain of isolated populations in small patches that on the whole remained connected. Even in very cold conditions, no continuous connectivity of core areas seems to have been possible between the Eastern Cordillera and Cordillera de Mérida, or the region of Sierra de Perijá. Towards the south of the Eastern Cordillera a low-elevation barrier was possibly crossed at 1900 m asl forming a brief bridge suitable for páramo habitat into the Macizo Colombiano of the Central Cordillera.

**Figure 4.**
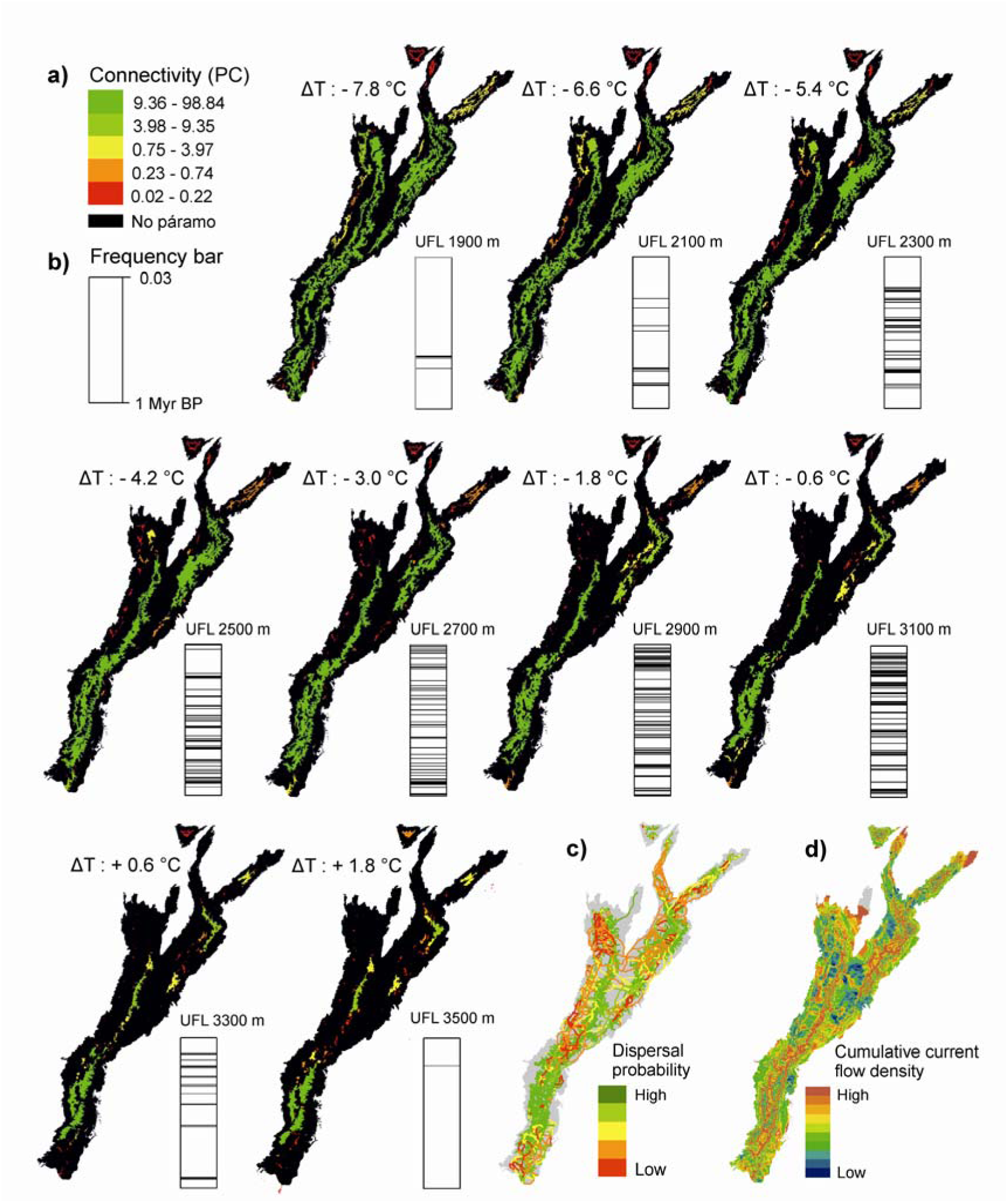
Páramo connectivity at different upper forest line (UFL) elevations. a) Probability of connectivity metric (PC; distance = 10 km, p = 0.5) (Saura & Rubio, 2010) calculated for all páramos larger than 1 km^2^. Maps are plotted with natural-breaks classification. Temperature at 2550 m elevation are relative to the present. b) Frequency bar indicates when the corresponding UFL elevation occurred during the last 1 Myr. Dispersal pathway analysis among páramos using c) Least cost pathways and d) Circuit model expressed in cumulative current flow density (McRae et al., 2008). Areas with high dispersal probability (c) and high current flow (d) indicate frequent and highly probable corridors during the last 1 Myr (weighted by frequency and duration). See Appendices 4 and 5 for all maps and frequencies.

The reconstruction of putative corridors shows a complex spatial pattern through the mountainous landscapes of the Northern Andes (**Fig. 4c,d**). The long ridge of the Central Cordillera forms the starting point of numerous corridors to the páramos in the Western Cordillera. The Eastern Cordillera shows a complex internal pattern of corridors, where there are neither strong corridors towards Sierra de Perijá in the North, nor towards the Cordillera de Mérida, while a high concentration of corridors is found between the large páramos complexes in the Eastern Cordillera (Páramos of Boyacá and Cundinamarca, **Fig. 1**). In the Ecuadorian Cordillera a more lateral pattern of high/low potential corridors is observed following the intra-Andean valleys and peaks within this mountain range. Corridors to the southernmost páramos of Ecuador as also the northernmost páramos of the Western Cordillera are weak and occurred infrequent during the last million years, shown by the thin lines.

### 3.3 Flickering connectivity systems

Páramo connectivity through time shows a highly variable pattern (**Fig. 5.a**) introduced by Flantua & Hooghiemstra (2018) as a flickering connectivity system (**see visualization in Appendix S6**). We find support for the hypothesis that this system with fluctuating, highly variable connectivity in spatial and temporal dimension is unique for each mountain range of the Northern Andes (**Fig. 1**). For example, changes in connectivity within the Ecuador Cordillera are substantial but the system ‘flickers’ around a high average when compared to other mountain ranges. The flickering connectivity systems within the Eastern and Central Cordillera are surprisingly similar, though the peaks of connectivity during glacial periods and the dips of connectivity during interglacials are more extreme in the former (**Fig. 5a**). The Western Cordillera is a larger mountain range than the Cordillera of Mérida and the SNSM (**Table S.1**), and its variation of connectivity has been correspondingly larger (**Fig. 5b**) but with the lowest occurrence of connectivity compared to the other mountain ranges (**Fig. 5a**). Considering only the frequency in the distribution of data (**Fig. 5b**), the Ecuadorian Cordillera and the SNSM stand out for their relatively small within-mountain range variation in connectivity, compared to the Eastern and Central Cordillera (similar patterns) and the Western Cordillera.

**Figure 5.**
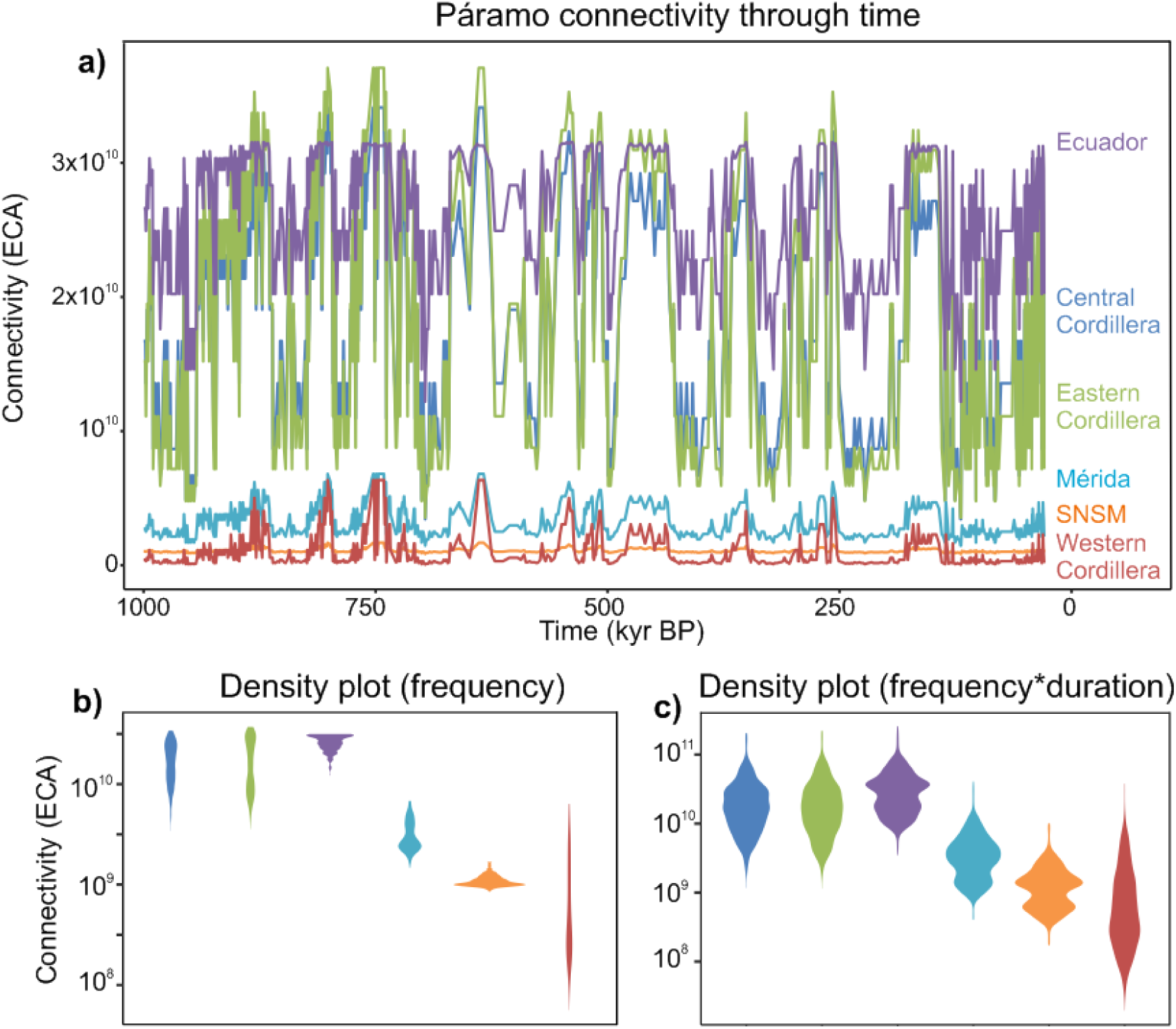
The ‘Flickering Connectivity System’ of the Northern Andes. a) Páramo connectivity (ECA) through time (3000-30 kyr BP) for each cordillera. b) Violin plot showing the distribution of the data and its probability density. Plot only considers how often certain connectivity occurred, not how long it lasted. c) Violin plot showing the distribution of the data and its probability density multiplied by how long connectivity persisted.

When frequencies of connectivity are weighted by the amount of time that connectivity lasted two main patterns emerge (**Fig. 5c**). The first is shared by the Western, Central, and Eastern Cordilleras, which all display an elongated pattern where the highest values are around a centroid, resembling a ‘humming top’ or, as Elsen & Tingley (2015) recognized in mountain hypsographies, a ‘diamond’ shape. Ecuadorian Cordilleras, Cordillera de Mérida and SNSM instead reveal a different pattern with a narrower centroid that widens towards the upper and lower section, resembling an ‘hourglass’ shape. Here, the Ecuadorian Cordillera and SNSM show a surprising similarity though at different connectivity ranges. The Central and Eastern Cordilleras are strikingly similar overall.

## 4. DISCUSSION

### 4.1 Variable degrees of past connectivity of different mountain ranges

Although currently isolated, evolutionary radiations and the assembly of the páramo ecosystem formed during times when the páramos were flickering in and out of connectivity (**Fig. 5b**.) The concept of ‘mountain fingerprints’ (Flantua & Hooghiemstra 2018) proposes that the region’s complex topography would have meant that páramos in different mountain regions would have fragmented and connected at different periods of time and with different rates and frequencies (as summarized in **Fig. 1**). This means that in some mountain ranges the páramos are a mix of somewhat even occurrence of connectivity and fragmentation events through time (**Fig. 1b**, here representative of the Eastern Cordillera), or could have been dominantly fragmented (**Fig. 1a**, e.g. Western Cordillera), or more connected (**Fig. 1c**, e.g. Ecuadorian Cordilleras). These regional differences in the temporal and spatial variation in past páramo connectivity (**Figs 4** and 6) are likely to have resulted not only in regional differences in biogeographical patterns through time, but also varying ecological and evolutionary processes. We therefore propose that the data we present can be used to test hypotheses of the drivers of species richness, endemism and degrees of Pleistocene diversification in the Northern Andes, and likewise are applicable to other mountain regions around the world.

### 4.2 Evolutionary implications of the flickering connectivity system

The dynamic history of the páramos elucidated by the flickering connectivity system can provide three important insights in terms of evolutionary processes. First of all, the regional differences in past páramo connectivity - the mountain fingerprint – support temporally and spatially discordant phylogeographic patterns (Pennington et al., 2010; Massatti & Knowles, 2014; Papadopoulou & Knowles, 2015; 2016). This means that the timing of diversification in the different mountain regions would not be expected to have occurred synchronously, even if all phylogenetic studies on páramo species could overcome current issues in techniques, spatial resolution and time-calibration points (Rull, 2011). Secondly, diversification rates might differ along the elevational gradient and this might be the rule rather than the exception. Elevational differences in surface availability and connectivity are likely to influence at what elevation the strongest phylogeographic processes will occur, and these processes are thus expected to differ between mountain systems resulting in elevational differences of diversification (see for instance Kropf, Kadereit, & Comes, 2003; Lagomarsino, Condamine, Antonelli, Mulch, & Davis, 2016). And thirdly, the flickering connectivity system, which is expected to cause isolation followed by connectivity of populations, is expected to cause pulses of diversification (Knowles, 2000), possibly resulting in series of sub-radiations in the páramos. Where isolation resulted in allopatric, *in situ* speciation, connectivity triggered diversification through dispersal and settlement in new areas (“dispersification”, Moore & Donoghue, 2007), and hybridization of differentiated taxa from previously isolated populations (Petit et al., 2003; Grant, 2014). Phylogenetic studies are increasingly supportive of the important role of gene flow, dispersification and hybridization, alongside periods of isolation, in driving (explosive) diversification in mountains (e.g. Knowles & Massatti, 2017; Hazzi, Moreno, Ortiz-Movliav, & Palacio, 2018), as well as in other systems such as tropical rainforests (e.g. Onstein et al., 2017) and islands (e.g. Ali & Aitchison (2014). In the páramos, examples originate from studies on birds (Quintero & Jetz, 2018; Cadena et al., 2016) and plants, such as *Neobartsia* (Uribe-Convers & Tank, 2015), *Lupinus* (Hughes & Eastwood, 2006; Nevado, Contreras-Ortiz, Hughes, & Filatov, 2018; Contreras-Ortiz, Atchison, Hughes, & Madriňán, 2018), *Loricaria* (Kolář et al., 2016), *Espeletia* (Diazgranados, 2012; Diazgranados & Barber, 2017; Pouchon et al., 2018) and *Hypericum* (Nürk, Scheriau, & Madriñán, 2013), supporting the strong relationship between changing degrees of connectivity and radiations (Flantua & Hooghiemstra, 2018). Interestingly, the Funza09 pollen record shows a clear shift in the rhythm of climate change around the mid-Pleistocene transition (ca. 0.9 Ma) after which 100 kyr cycles with high amplitudes started to dominate the climate, overruling the lower-amplitude 41-kyr cycles that continued in the background. Strikingly, changes in speciation rates of high elevation birds (Weir, 2006) and the Espeletiinae in the Cordillera de Mérida (Pouchon et al., 2018), echo the mid-Pleistocene transition by an acceleration of diversification during the last 1 Myr suggesting a close link between the intensity of the flickering connectivity system and radiations.

### 4.3 Limitations and model assumptions

Inherent to any model in mountains and concerning connectivity are simplifications of the temporal and spatial complexity of the real world. For instance, the UFL is asymmetric on wet and dry mountain slopes (e.g. Cleef, 1981), the current elevation of the UFL shows a range of variation of ca. 200 m (incidentally to 300 m), surface processes have changed topography on a million years time scale (Herman et al., 2013; Antonelli et al., 2018), the elevational temperature gradient (lapse rate) seems higher during glacial conditions than at present (Wille, Hooghiemstra, Behling, van der Borg, & Negret, 2001; Loomis et al., 2017), and the current subdivision of páramo vegetation into a 300 m: 600 m: 200 m interval for shrubpáramo, grasspáramo, and superpáramo, respectively, is subject to change (Van der Hammen, 1981; Hooghiemstra, 1984), potentially related to changing atmospheric pCO_2_ levels (Boom, Mora, Cleef, & Hooghiemstra, 2001; Boom, Marchant, Hooghiemstra, & Sinninghe Damsté, 2002). We estimate the potential impact of these factors on the estimated connectivity of little significance in determining the overall patterns observed in the flickering connectivity systems.

Any study concerning connectivity also uses a number of assumptions on the probability of dispersal through the landscape. Here, we used a generalized PC value of 0.5 at 10 km to estimate the degree of connectivity. However, species traits, life histories and dispersal capacities may strongly influence dispersal distances (Onstein et al., 2017), and thus influence the probability of connectivity between populations. Implementing taxon-specific traits when calculating the landscape resistance grid (see Methodology) may thus improve the accuracy of the connectivity estimates. Also, family or taxa specific connectivity maps could uncover why certain plant genera do not show any evolutionary diversification during the Pleistocene, e.g. Distichia (Juncaceae; Colin Hughes, personal comm.) and Arcytophyllum (Rubiaceae; Madriñán, Cortés, & Richardson, 2013). Additionally, *a priori* “hard” barriers can be imposed to emphasize areas where habitat connectivity is unlikely to have occurred (see for instance how we maintained SNSM isolated from the other mountain ranges). Defining these barriers *a priori* is not indispensable, though, as the connectivity analysis hints at strong dispersal restrictions when resistance values of corridors are high and indicative of highly constrained dispersal. In the Northern Andes, this is shown by the multiple single line corridors between the Central and Eastern Cordillera, confirmed by the lack of gene flow between plant populations of these regions (Jabaily & Sytsma, 2013; Diazgranados & Barber, 2017; Contreras-Ortiz et al., 2018). This example illustrates the added value of integrating different lines of evidence (e.g. genetic, fossil, paleoclimate) in a spatial and temporal context to understand the biogeographical patterns observed in phylogeographic studies.

### 4.4 Future research

Our spatio-temporal estimates of past connectivity lay a foundation for further research on elucidating the causal mechanisms of mountain diversifications. Models of past connectivity (**Figs. 4** and **5**), when combined with phylogeographic data, could help reveal the role of interspecific gene flow and allopatric speciation in driving radiations in the high Andes (Nevado et al., 2018) and contribute to a better understanding of the relative importance of geography vs adaptive radiation that underpin Andean diversifications (Contreras-Ortiz et al., 2018). In such a complex system it may also be useful to pay attention to commonalities. For example, when considering both frequency and duration, our data show that two connectivity patterns emerge (i.e. hourglass vs. non hourglass; **Fig. 5.c**). Research could explore if cordilleras with shared connectivity patterns also share phylogenetic histories and contemporary (endemic) species’ biogeographies to test for universal mechanisms that have shaped present day alpine biomes. This would be especially useful if used in conjunction with information on the reproductive life histories, growth and dispersal capacities of specific taxa.

Finally, past patterns of connectivity are critical to interpret biogeographical patterns of currently isolated or fragmented systems in a wide variety of terrestrial ecosystems including mountains (Flantua & Hooghiemstra, 2018), islands (e.g. Simpson, 1974; Weigelt, Steinbauer, Cabral, & Kreft, 2016; Norder et al., 2018), fresh water systems (e.g. Dias et al., 2014), rainforests (e.g. Graham, Moritz, & Williams, 2006), and grasslands (e.g. Lindborg & Eriksson, 2004; Münzbergová et al., 2013), and marine coastal ecosystems (Hoeksema, 2007) that similarly experienced major spatial changes during rapid sea-level fluctuations over the Pleistocene. The approach developed here, to quantify historical connectivity, can therefore provide a platform for interpreting contemporary biogeographies and past drivers of diversification in a wide array of both marine and terrestrial ecosystems where available space has been altered by climatic fluctuations. We postulate that quantifying flickering connectivity systems will facilitate a much more detailed and much needed quantitative basis to compare diversity patterns across the mountain regions of the world.

## 5. CONCLUSIONS

We present a pollen-based biogeographical model for the páramos biome spanning the northern Andes (Venezuela, Colombia and Ecuador) over the last 1 Myr. Our models suggest substantial temperature oscillations where extreme temperature lows were ca. 8°C cooler than today, causing a lowering of the UFL of ca. 1600 vertical meters. These extreme cool events were however rare and during glacial periods most of the time cool stadials and interstadials prevailed. Our analysis shows that páramos on all mountain ranges underwent frequent alterations between fragmented and connected configurations, but the estimated degrees and amount of connectivity varied among mountain ranges. Most páramos expanded during glacial periods even though extensive glaciers were present. To a large extent the current páramo distribution was replaced by glaciers, implicating a substantial range size change of populations and a highly dynamic system during Pleistocene times.

In light of Von Humboldt’s work of relevance of different topographies for mountain biota, we show that topography and climate change together dictated páramo connectivity through time with high spatial variability. The interplay of the topographic and paleoclimatic conditions created a unique pattern of connecting and fragmenting páramo patches through time, here described as the flickering connectivity system. Our spatially explicit model quantifies the complexity of mountain biome dynamics during climate oscillations, in terms of the degree, frequency and duration of past connectivity of alpine biome mountains (**Figs 4** and **5**) and can be applied to other mountain regions. Our connectivity estimates can contribute to answering long-standing questions on the drivers of evolutionary diversification in phylogenetic and phylogeographic studies, and enrich our understanding of the biogeographic history of mountain ecosystems more generally.

> *“There the different climates are ranged the one above the other, stage by stage, like the vegetable zones, whose succession they limit; and there the observer may readily trace the laws that regulate the diminution of heat, as they stand indelibly inscribed on the rocky walls and abrupt declivities of the Cordilleras”* (Von Humboldt, 1877 (1845), I, p 46).

## ACKNOWLEDGEMENTS

This work was part of SGAF’s doctoral thesis funded by Netherlands Organization for Scientific Research (NWO, grant 2012/13248/ALW to HH.). The Hugo de Vries foundation is acknowledged for financially supporting multiple grant proposals during the project including the development of the visualization accompanying this paper. The Sistema Nacional de Investigadores (SNI) de SENACYT supported AO. REO acknowledges the support of the German Centre for Integrative Biodiversity Research (iDiv) Halle-Jena-Leipzig funded by the Deutsche Forschungsgemeinschaft (DFG, German Research Foundation)— FZT 118. Carina Hoorn and Daniel Kissling are thanked for the educational environment shaped by the parallel paper on mountain diversity (Antonelli et al., 2018). Colin Hughes is thanked for comments on previous versions of this paper. We thank Mauricio Bermúdez for help with the geological delimitation of the Northern Andes and Francisco Cuesta for facilitating the map by Josse et al. (2009).

## BIOSKETCHES

**Suzette Flantua** has a background in paleoecology, biogeography, landscape ecology and spatial analyses, and enjoys integrating them all. She is interested in a wide range of topics from the Miocene to the present, from islands to mountains, to understand contemporary patterns of species richness and endemism.

**Henry Hooghiemstra** is a terrestrial and marine tropical palynologist working on time-scales from the full Quaternary to the Anthropocene. His research focuses on a wide variety of biomes in Central and South America, Saharan and East Africa and in Mauritius.

**Aaron O’Dea** is a marine paleobiologist and uses the marine fossil record in Tropical America to explore drivers of macroevolution in the seas, and takes cores on coral reefs from French Polynesia to the Dominican Republic to reconstruct how reefs changed over millennia with the aim of improving their future resilience.

**Renske E. Onstein** is an evolutionary ecologist who enjoys collecting (and eating) tropical megafaunal fruits, e.g. on Borneo and Madagascar, while studying how fruit functional traits interact with frugivores to affect diversification dynamics. She is generally interested in the broad-scale distribution and diversification of functional and taxonomic diversity of flowering plants.

## APPENDICES

Additional Supporting Information may be found in the online version of this article:

**Appendix 1 |** Surface areas, elevational ranges and hypsographies of the Northern Andes

**Appendix 2 |** Background on the páramo alpine biome

**Appendix 3 |** Methodology underlying the use of fossil pollen data to reconstruct the upper forest line changes

**Appendix 4 |** Degree of connectivity of páramos at all UFL elevations during the last 1 Myr.

**Appendix 5 |** Frequency analysis of all UFL elevations during the last 1 Myr.

**Appendix 6 | Visualization of the flickering connectivity system in the Northern Andes.** Artwork by Catalina Giraldo Pastrana in collaboration with Suzette G.A. Flantua and Henry Hooghiemstra.

**Appendix 7 |** Further suggestions for future work.

## AUTHOR CONTRIBUTIONS

S.G.A.F. and H.H. conceived the ideas. H.H. provided the AP% of the Funza09 dataset. S.G.A.F. performed the spatial analyses. S.G.A.F and H.H. led the writing and figure design with critical contributions by A.O. and R.E.O. All authors contributed to versions of the manuscript and revisions.

