## Appendix S1 for "The flickering connectivity system of the north Andean páramos"

**SUPPORTING INFORMATION**

### Appendix S1. Surface areas, elevational ranges and hypsographies of the Northern Andes

The highest peaks of the Northern Andes are located in Ecuador (ca. 6300 m asl) and Sierra Nevada de Santa Marta (SNSM; ca. 5800 m asl). In Colombia, three parallel positioned cordilleras cross in north-south and northeast-southwest direction the region, separated by deep inter-Andean valleys, of which the highest is the Central Cordillera. They join in Ecuador to form a high elevation mountain block with numerous volcanos throughout the region as often displayed in the work by Von Humboldt & Bonpland (1805; 1807). The largest cordillera is the Eastern Cordillera, followed by the Central Cordillera and the Ecuadorian Andes (**Table S1.1**). The SNSM is the smallest but highest in Colombia, forming an isolated mountain range just of the Caribbean coastline.

| Mountain range | Total surface area ($km^2$) | Elevational range (m asl) | Hypsographic shape (Elsen & Tingley, 2015) | Hypsographic shape (this study) |
|---|---|---|---|---|
| Northern Andes | 448.000 | 13-6300 | Hourglass (total Andes) | Pyramid |
| Sierra Nevada de Santa Marta (SNSM) | 9190 | 430-5700 | Pyramid (included in Eastern Cordillera) | Pyramid |
| Cordillera de Mérida | 30.340 | 80-5000 | Pyramid | Diamond |
| Eastern Cordillera | 130.630 | 106-5400 | Pyramid | Hourglass/Pyramid (with high elevation plateau) |
| Central Cordillera | 122.000 | 81-5300 | Diamond (joined with Ecuador) | Pyramid |
| Western Cordillera | 48.100 | 13-4100 | Pyramid | Diamond |
| Ecuadorian Cordilleras | 107.800 | 25-6300 | Diamond | Hourglass/Diamond |

**Table S1.1 |** Surface availability and hypsographic shape for each mountain range of the Northern Andes.

To understand elevational availability of surface areas in mountains, hypsographic curves provide estimates of elevational differences (See for a global analysis of mountains Elsen & Tingley, 2015). To calculate these curves for each cordillera, we first delimitated the Northern Andes by using the boundary defined by Josse et al. (2009). Then we added the SNSM by a 500 m asl isoline as the previous study included all but this mountain range. To define the boundaries between the three different cordilleras of Colombia, and between the Central Cordillera in Colombia and the Ecuadorian Cordillera, we identified the main geological faults following Bermúdez, Van der Beek & Bernet (2013), Baldock (1982) and Ramos & Alemán (2000), and then provided additional detail to the cordillera boundaries by defining the deepest sections of the main canyons using ESRI ArcHydro tool (ESRI, 2014).

Von Humboldt, A., & Bonpland, A. (1805). *Essai sur la Géographie des Plantes; Accompagné d'un Tableau Physique des Régions Équinoxiales*. Paris: Chez Levrault, Schoell.

Von Humboldt, A., & Bonpland, A. (1807). *Geographie der Pflanzen in den Tropenlandern*. Paris.
