## Appendix S2 for "The flickering connectivity system of the north Andean páramos"

**SUPPORTING INFORMATION**

### Appendix S2. The páramo alpine biome

Plant species richness in the páramos estimates range from ca. 3500 (Sklenář, Dušková, & Balslev, 2011) to ca. 3700, much of which are in the families Asteraceae, Orchidaceae and Poaceae (Rangel, 2015). The most striking taxonomic difference among the páramos of the Northern Andes is the presence of the emblematic stem rosette of the Espeletiinae in Venezuela and Colombia, but which is absent in most páramos in Ecuador. Páramos along the east side of the Northern Andes (facing the Amazon basin) are also generally more humid than those along the west side (facing the Pacific). The páramos of the Eastern Cordillera of Colombia currently cover the largest surface area (ca. 15,800 $km^2$) compared to those of the Central Cordillera (ca. 8200 $km^2$) and the Western Cordillera of Colombia (ca. 690 $km^2$).

Traces of extensive glaciers during glacial times are found throughout the Northern Andes (Helmens, 1990; Schubert & Clapperton, 1990; Clapperton, 1993; Braun & Bezada, 2013), having shaped and carved the high elevation landscape of the páramos during a significant part of the last million years. In current times, the largest ice covers are found around the area of the páramos of Boyacá, Sierra Nevada El Cocuy and Chimborazo (**Fig. S2.1)**, exposed to the moisture supply from the Amazon basin. Most Andean glaciers, however, are small remnants of what were once massive glaciers covering most, if not all, mountain ridges, but are now either rapidly decreasing or have disappeared in the last decades (IDEAM, 2012); Braun & Bezada, 2013).

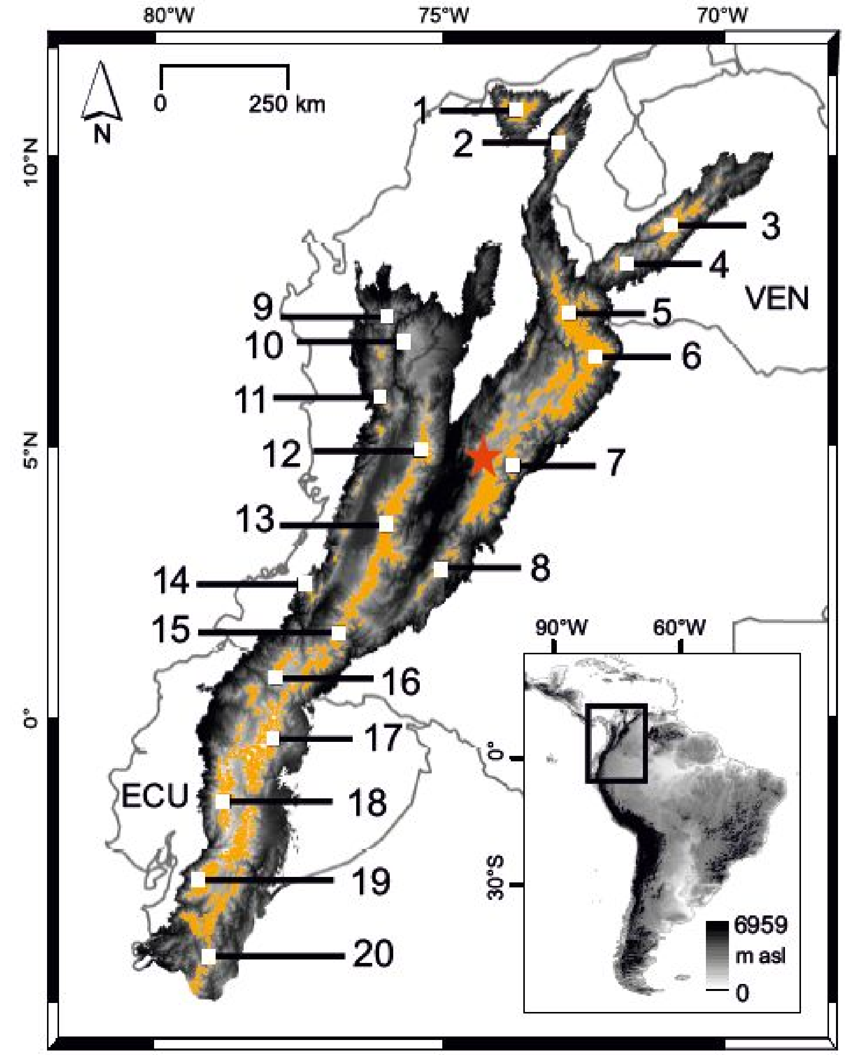


**Figure S2.1 | The Northern Andes showing the current distribution of páramos**. Main páramos are: 1. Sierra Nevada de Santa Marta, 2. Sierra de Perijá, 3. Sierra Nevada de Mérida and Santa Domingo, 4. Páramo del Batallon, 5. Páramos de los Santanderes, 6. Páramos de Boyacá, 7. Páramos de Cundinamarca, 8. Páramos Los Picachos and Miraflores, 9. Páramo Paramillo, 10. Páramo Belmira, 11. Páramos Frontino-Tatamá, 12. Páramos Viejo Caldas-Tolima, 13. Páramos Valle-Tolima, 14. Páramos del Duende-Cerro Plateado, 15. Páramos Macizo Colombiano, 16. Páramos del Antisana, 17. Páramos de Cayambe, 18. Páramos de Tabacundo, 19. Páramos de Chimborazo, 20. Páramos de Cuenca. Red star indicates location of fossil pollen record of Funza09 (Torres, Hooghiemstra, Lourens, & Tzedakis, 2013). VEN: Venezuela; COL: Colombia; ECU: Ecuador. Northern Andes limits adapted from Josse et al. (2009) and páramos as defined by Sarmiento Pinzón, Cadena Vargas, Sarmiento Giraldo, & Zapata Jiménez (2013) and IAvH (2012).
