## Appendix S3 for "The flickering connectivity system of the north Andean páramos"

**SUPPORTING INFORMATION**

### Appendix S3 | Methodology underlying the use of fossil pollen data to reconstruct the upper forest line changes

Long and continuous records of past climate change of the Pleistocene, in mountains in particular, are rare. Due to a unique combination of subsidence and sediment infill, the intramontane basin of Bogotá, Colombia (4.83°N, 75.2°W; 2550 m asl) accumulated sediments during most of the Pleistocene (Torres, Hooghiemstra, Lourens, & Tzedakis, 2013). This interval starts with climate change driven by obliquity (41 kyr rhythm) and from ca. 0.9 Ma onwards the eccentricity (100 kyr) driven frequency overrules the obliquity signal, showing the change in glacial-interglacial dynamics after the mid-Pleistocene transition (1-0.9 Ma; Imbrie et al., 1984; Lisiecki & Raymo, 2005).

The UFL is the transition from the upper montane forest to the páramo, i.e., the uppermost contour of closed forest (Bakker, Moscol Olivera, & Hooghiemstra, 2008), which coincides with the ca. 9.5 °C mean annual temperature (MAT) isotherm (Hooghiemstra, 1984; Groot et al., 2011; Hooghiemstra et al., 2012). Following the reconstructions by Hooghiemstra (1984) and Groot et al. (2011), we assumed that during the last 0.4 Myr that Oak (*Quercus*) was present the empirical level of 40% AP concurs with having the UFL at the level of the Bogotá basin (2550 m asl). For the period after the arrival of Alder (*Alnus*; 1.01 Ma) and before the arrival of Oak (0.43 Ma) a 35% AP level is used. We applied the empirical relationship of 5% increments in AP% as reflecting steps of 100 m UFL displacement following Hooghiemstra (1984). We used a lapse rate of 0.6°C/100 m and an offset of -5.8°C to calculate fluctuations in temperature through time.

In our model, the páramos occupied an elevational range of 1200 m during the Pleistocene (Hooghiemstra, 1984; Groot et al., 2011; Bogotá-Angel et al., 2011; Bogotá-A., Hooghiemstra, & Berrio, 2016; Torres et al. 2013), consisting of subpáramo (ca. 300 m; dominated by shrub), grasspáramo (ca. 700 m; dominated by herbs), and súperparamo where vegetation is limited to the lowermost c. 200 m of the 600 m wide belt (Cleef, 1981). The elevational range above the páramos was assigned to being glaciers representing the extensive ice sheets present in the Northern Andes during colder and humid periods (Helmens, 1990; Schubert & Vivas, 1993; IDEAM, 2012). Páramos and glaciers show an asymmetrical zonation of the vegetation belts between the dry and wet side of the mountain (Cleef, 1981; Schubert & Clapperton, 1990), and also regional differences among cordilleras are observed (Moreno, Andrade, & Ruíz-Contreras, 2017). For instance, the UFL is often located at higher elevations when the mountain range is larger or higher, or along the atmospherically humid side of the mountain (but see the Eastern Cordillera; Van der Hammen & Cleef, 1986). We simplified these regional differences in our models by using the same elevational gradients for all mountain ranges, as these regional differences are unknown for the past and humans have modified the natural limit of the UFL through high elevation agriculture and reforestation (Bakker et al., 2008).
