## Appendix S4 for "The flickering connectivity system of the north Andean páramos"

### SUPPORTING INFORMATION

### Appendix 4 | Degree of connectivity of páramos at all UFL elevations during the last 1 Myr

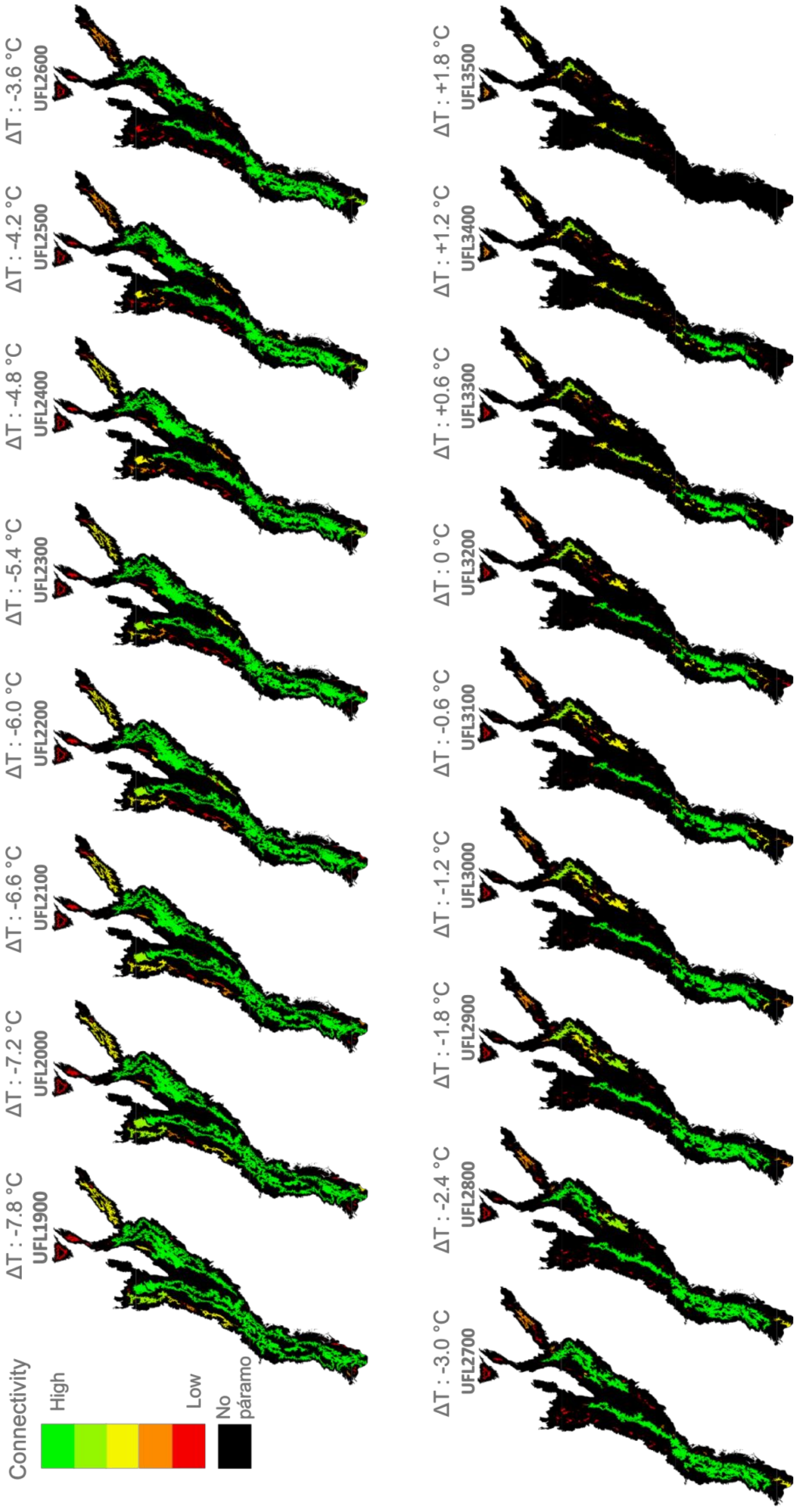
