## Appendix S5 for "The flickering connectivity system of the north Andean páramos"

### SUPPORTING INFORMATION

### Appendix 5 | Frequency analysis of all UFL elevations during the last 1 Myr

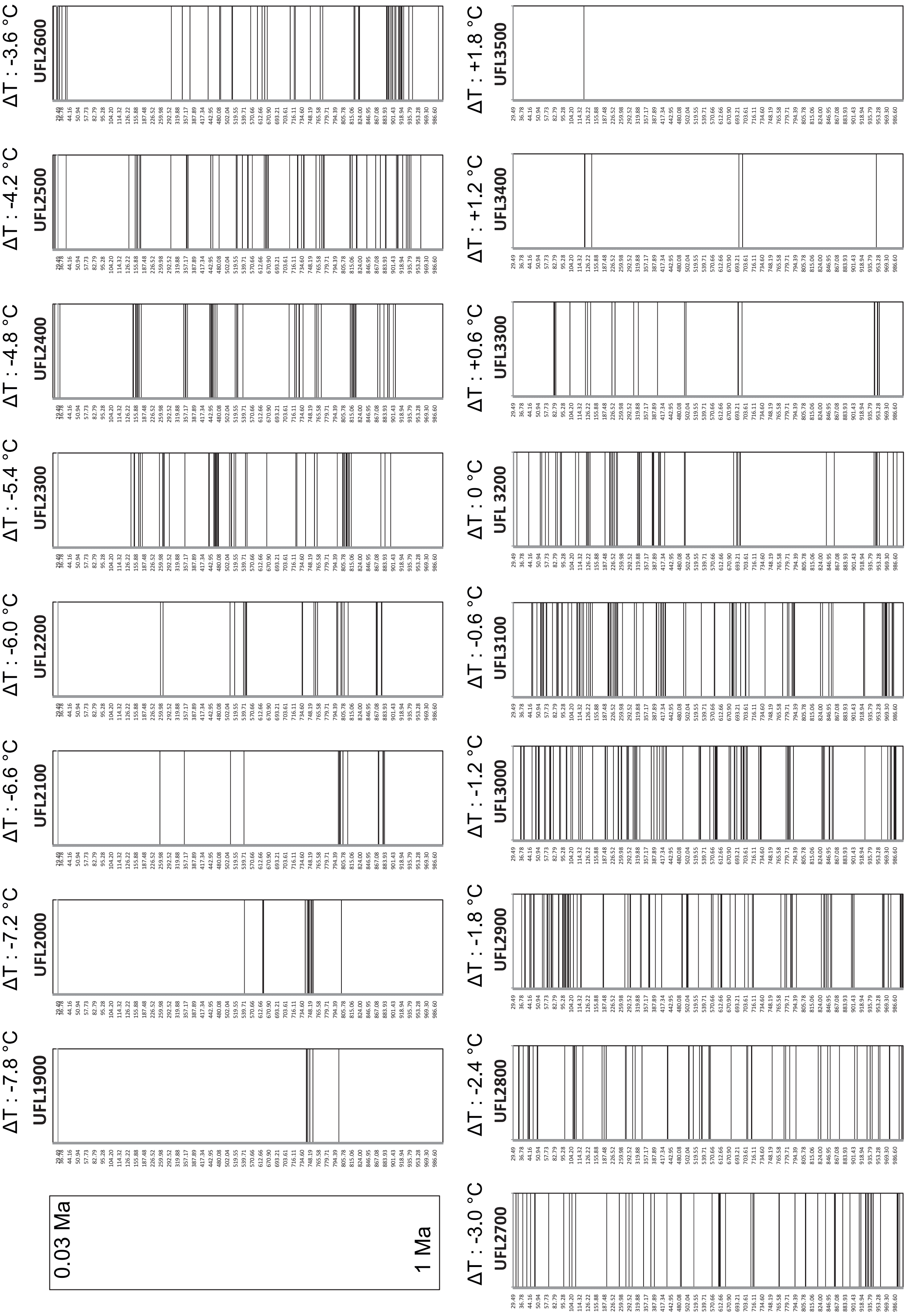
