## Appendix S6 for "The flickering connectivity system of the north Andean páramos"

**SUPPORTING INFORMATION**

### Appendix 6 | Visualization of the flickering connectivity system in the Northern Andes.

Artwork by Catalina Giraldo Pastrana in collaboration with Suzette G.A. Flantua and Henry Hooghiemstra.

The flickering connectivity system has now been visualized in the form of a 3D animation and photography film. The visualization was created by Catalina Giraldo as student of Master in Arts in Media Design and Communication at the Piet Zwart Institute, Rotterdam University of Applied Sciences, and in collaboration with Suzette G.A. Flantua and Henry Hooghiemstra at IBED, University of Amsterdam.

Please follow the following link to access the video (ca. 1.1GB size):

https://figshare.com/s/a288ff6903462a8fb7af

This is a temporal link. Permanent link will be made available upon final publication and the following doi will become active: 10.6084/m9.figshare.7408643
