## Appendix S7 for "The flickering connectivity system of the north Andean páramos"

**SUPPORTING INFORMATION**

### Appendix S7 | Further suggestions for future work

Future work would ideally incorporate regional differences in paleoclimate as our model was based on only one paleotemperature curve. However, though fossil pollen records in the Andes are numerous, long fossil pollen records similar to Funza-09 are currently not available (Flantua et al., 2015). While paleoclimate reconstructions based on computer climate models are in rapid development, and could be of use for exploring the generality of patterns observed here, they are currently restricted to selected periods in time: the Last Glacial Maximum, last interglacial and the mid-Holocene – but see Holden et al. (2018) and Rangel et al. (2018) – and show substantial underestimation of glacial cooling in mountainous areas (Bush & Philander, 1999; Loomis et al., 2017). We recommend the acquisition of more long pollen records, not only to improve our models, but also provide better understanding of climatic and biotic changes in this biologically-important part of the world.

Exceptionally long paleorecords also provide key insights into the degree of resilience of alpine biomes under strongly changing climates. Models often predict massive mountain top extirpations as taxa 'run out of space' and thus extinctions seem inevitable during warmer temperatures than the present. The páramos endured massive habitat replacement and substantial fragmentation, but nevertheless are famously known for their high number of plant radiations (see overview by Hughes & Atchison, 2015) and rapid diversification of high elevation birds (Weir, 2006; Quintero & Jetz, 2018). Interestingly, there are no observed extinctions of plant taxa in fossil pollen records that cover several glacial-interglacial cycles (Torres, Hooghiemstra, Lourens, & Tzedakis, 2013; Groot et al., 2011; Bogotá-Angel et al., 2011), a phenomenon previously described to be the "Quaternary conundrum" (Botkin et al., 2007). Even though pollen data do not reach species level, pollen records do represent a representative cross section of the vegetation observing both common and rare species, and thus complete disappearances should be observable (Colinvaux & Schofield, 1976). How then has this exceptionally rich biome with numerous endemics flourished under such dynamic and often highly fragmented conditions, and what can be learned for improving predictions of the future of these and other geographically-limited habitats under climate change scenarios? Forecasting models likely need further fine-tuning to include other genetic and ecological mechanisms that come into play when páramos are reduced to mountain tops, and strategies that reduce rates of extinctions become evident (Botkin et al., 2007). Another possible explanation is that the vegetation belt structure, so elegantly drawn by (Von Humboldt & Bonpland, 1805), loses its arrangement towards the top of the mountain and populations reassemble into different configurations, which buffers against extinction, possibly a mosaic patterning, but further research from different mountain systems is needed to test such hypothesis.
